# Non-parametric physiological classification of retinal ganglion cells in the mouse retina

**DOI:** 10.1101/407635

**Authors:** Jonathan Jouty, Gerrit Hilgen, Evelyne Sernagor, Matthias H. Hennig

## Abstract

Retinal ganglion cells, the sole output neurons of the retina, exhibit surprising diversity. A recent study reported over 30 distinct types in the mouse retina, indicating that the processing of visual information is highly parallelised in the brain. The advent of high density multi-electrode arrays now enables recording from many hundreds to thousands of neurons from a single retina. Here we describe a method for the automatic classification of large-scale retinal recordings using a simple stimulus paradigm and a spike train distance measure as a clustering metric. We evaluate our approach using synthetic spike trains, and demonstrate that major known cell types are identified in high-density recording sessions from the mouse retina with around 1000 retinal ganglion cells. A comparison across different retinas reveals substantial variability between preparations, suggesting pooling data across retinas should be approached with caution. As a parameter-free method, our approach is broadly applicable for cellular physiological classification in all sensory modalities.

## Introduction

It is well established that the retina has multiple, functionally complementary populations of retinal ganglion cells (RGCs), which together transmit visual information to various central visual areas (1). Strikingly, the stratification of RGC dendrites in the inner plexiform layer predicts response polarity and kinetics extremely well. In addition, the retinotopic organisation of RGCs is such that each type independently covers the visual space through receptive field tiling as regular mosaics. The retina is among the first neural systems where a clear correspondence between morphology, physiology and function of different cell types has been established (2–9, 28), an organising principle that likely exists in other neural systems as well. The actual classification of RGCs typically requires a combination of measures of their cellular physiology, light responses, morphology and, more recently, their gene expression patterns and connectome (10–14).

Despite such well defined principles, in practice classification of RGC types has been challenging because it is difficult to implement all criteria in a single experiment. Classification is particularly challenging when attempted solely based on light responses. In previous studies, features were extracted from responses to a set of stimuli designed to reveal the main spatial and temporal receptive field properties, as well as specific properties such as direction selectivity (DS). Using this approach, Farrow and Masland (15) found 12 distinct types, which mirrors a similar number of morphological types identified through unsupervised clustering (6). Moreover, consistent patterns in spike trains have been shown to allow distinction between major RGC types (16). Numerous, more detailed studies have since refined and extended this picture, but without reaching a clear consensus. Recently Baden et al. (17), combining clustering, dimensionality reduction of peristimulus time histograms (PSTH) and other response criteria and morphological information, reported at least 39 distinct RGC types.

Previous studies thus appear to suggest that light responses alone do not contain sufficient information for reliable RGC classification unless a careful stimulus ensemble is designed to evoke optimal responses, in particular for specialised RGCs such as DS cells. Yet, if different RGC types have distinct cellular physiological properties, light responses, and systematic differences in their presynaptic input neurons, it may be possible to use a sufficiently rich stimulus to evoke responses to unmask their distinctive functional features. Here we propose that it is not necessary to use stimuli that optimally excite the receptive field of each RGC type. Instead, we expect that these cellular differences cause perhaps subtle, but detectable differences in the light responses even to non-optimal stimuli. The actual identity of each RGC type, including specific properties such as DS, can then be confirmed post-hoc using more specific stimuli.

A main caveat in previous studies was that RGC recordings from multiple preparations were pooled together to obtain a data set of sufficient size. Large scale, high density microelectrode arrays (MEAs) now make it possible to record large populations comprising thousands of cells from a single retina (18–21). This reduces contaminating effects of variability between animals and experimental conditions, and allows precise control of stimulation for all recorded neurons. Extending an idea first presented by Zeck and Masland (16), here we present a method for clustering RGCs based on spike distance measures, which is particularly suited for high density recordings. Its main advantage is that it is a parameter-free distance measure for clustering. We first validate the method using synthetically generated RGC spike trains. The results show that the methods using the parameter-free spike distances compare favourably to clusterings based on feature vectors, especially in the presence of low to medium levels of noise. On recorded RGC data, the method is able to distinguish many distinct RGC types, as confirmed by assessing response properties during stimulation that was not part of the data used for clustering. A comparison between different retinas shows that similar types can be identified, but that heterogeneity between preparations prevents the use of pooled data. Together our work suggests a new strategy for consistent identification of RGC types, and potentially of neurons in other sensory systems where appropriate stimulation paradigms can be designed.

## Methods

All code to reproduce the experimental data analysis presented in this paper, and an example data set is available at https://github.com/mhhennig/rgc-classification. This repository contains additional analysis, and hopefully provides a starting point for refinement and extension of the methods presented in this paper. We therefore encourage the reader to explore this resource, and to contribute to its improvement.

### Spike train distance based clustering

#### ISI distance

This measure is sensitive to dissimilarity in the inter-spike intervals (ISI) of two spike trains (22). The instantaneous ISI distance is the ratio between the ISIs, adjusted so that the distance is symmetric:

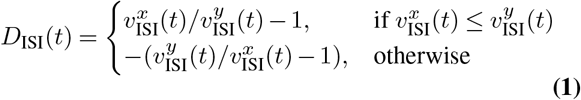

where 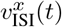 and 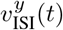 are the instantaneous inter-spike interval values for spike trains *x* and *y,* respectively.

#### SPIKE distance

The SPIKE distance compares the distances between the preceding and following spikes of the two spike trains (23, 24). First, the interval between the previous and following spike pairs are computed as

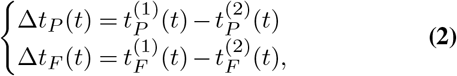

where 1 and 2 indicate the first and second spike train, and *P* denotes the spike pair before or at time *t*, and *F* the pair at or following *t*. Then a weighted average is computed as

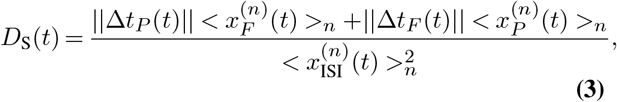

where 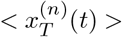 is the average of the intervals 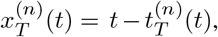 such that spikes closer together in time dominate the measure.

The ISI and SPIKE measures yield a quasi-instantaneous distance profile, which is averaged to obtain a single distance measure for each pair. These distances were computed using the open source package PYSPIKE ((25), version 0.5.3). Specifically, the pairwise distances between two units were determined by computing the pairwise distances of all trials of the same stimulus. The resulting distance matrix was then clustered with the hierarchical clustering algorithm as implemented in the Scipy library ((26), version 1.0.1), using the Ward variance minimization algorithm, as this gave the best results on ground truth data.

#### Synthetic RGC responses

##### Model

To evaluate the clustering quality, synthetic RGC spike trains emulating eight known base types, were generated using a linear-nonlinear-Poisson (LNP) model. The stimulus *s*(*t*) was similar to the chirp stimulus used in the MEA experiments (see below) (Fig. 1): 1.5 s darkness, followed by 2s full intensity and 2 s darkness; mid-point luminance ‘grey’ value for 2s; frequency modulation 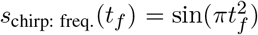 for 5s; 2s midpoint luminance; amplitude modulation ^*s*^_chirp: ampl._(*t_a_*) = 0.2*t* sin(3*πt_a_*) for 5 s; 2s midpoint luminance.

**Fig. 1.**
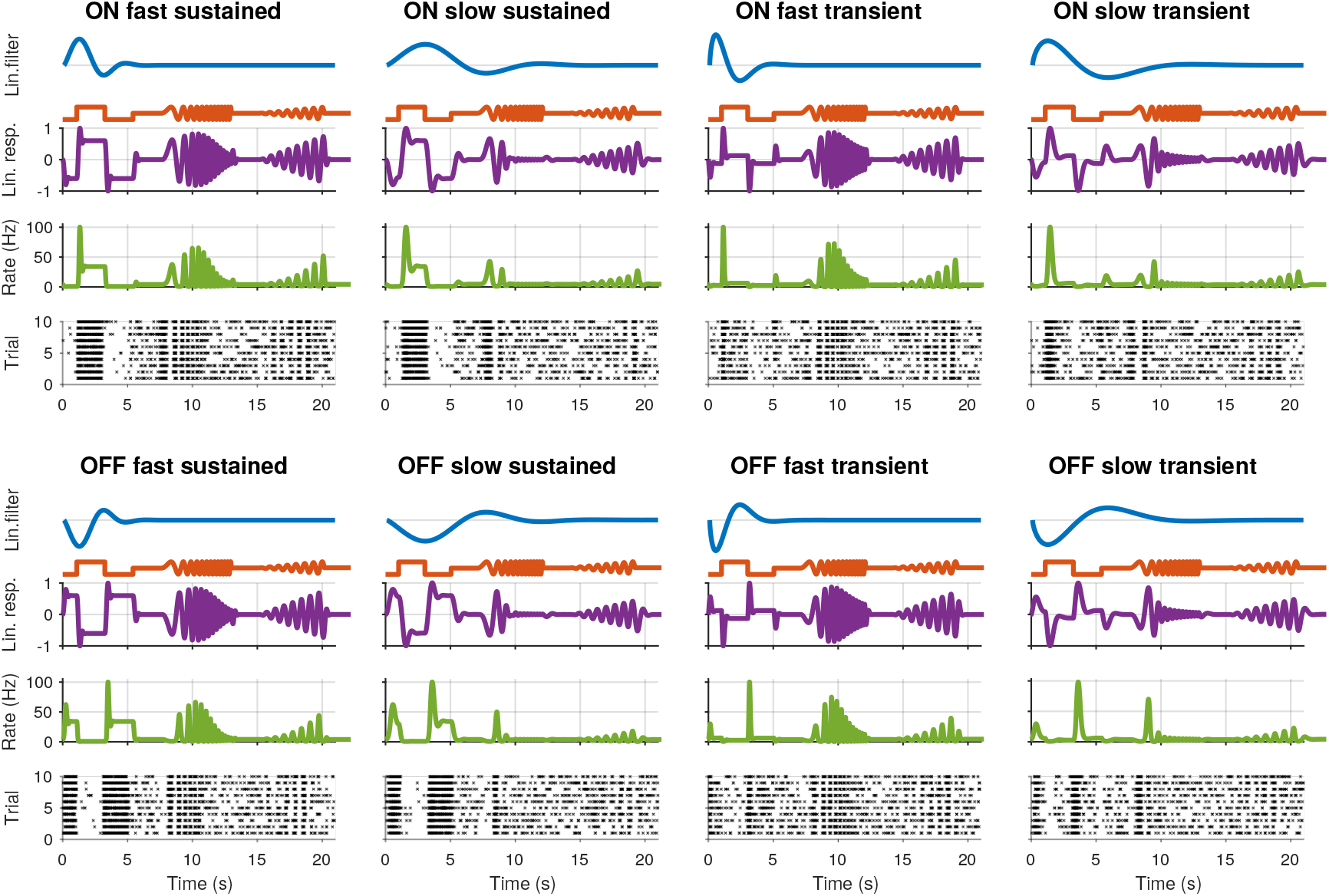
Synthetic retinal ganglion cell response models. For each of the eight cell types, rows from top to bottom: *(i)* the linear receptive field (RF) of the cell; *(ii)* the stimulus; *(iii)* the linear response of the cell, computed by convolving the RF with the stimulus; *(iv)* the spike rate obtained by passing the linear response through a non-linear function; and *(v)* the spike raster generated using a Poisson process.

We generated symmetric ON and OFF RGC types, with fast/slow and transient/sustained temporal characteristics, using Gabor functions with two parameters (27):

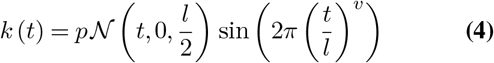

where 𝒩 is the Gaussian probability density function, and *p* the response polarity. This response is normalised in the range [*−*1, 1].

Here *l* is the “length” of the temporal receptive field, and modulates over how much the time the RF integrates the stimuli, and *v* is the “speed” that affect how quickly the cell responds to change in the stimuli intensity. A “long” AF thus averages out high frequency stimuli, whereas a “short” RF thus averages out high frequency stimuli, consistent with the main known distinguishing characteristics of RGC receptive field (12, 28). Note that these simulated RFs do not have spatial component, instead its effect during homogeneous illumination can be considered as parameterized by the temporal component importantly, the integral 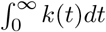 is constant for a fixed *v,* hence RFs with a different “length” but the same “speed” will have identical maximum linear response magnitudes, which allows testing the effect of response kinetics independent of response magnitude.

The cell’s RF filter 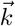 is convolved with the time-varying stimulus *s*(*t*) to yield the cell’s linear response. To simulate spikes, this response is passed through a non-linearnfunction *r*(*x*) which transforms the linear response into an instantaneous spike rate. We used a logistic sigmoid function:

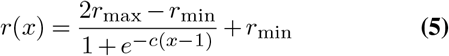

To match observations made in the recorded RGC spike trains, *r*_min_ was set to 0.5, and *r*_max_ to 100. By changing *c* and observing the effect on the different cell types, a value of 4.0 was found to result in the most balanced spike rate response across all types.

Finally, Poisson spike trains were generated in discrete time bins (*dt*) of 1 ms based on the instantaneous rate *P* (spike*|r*) = *r*(*t*) *× dt*.

##### Model parameter selection

We modelled eight cell types by combining the following parameters: “On” and “Off”: *p* = 1 and *p* = − 1; “Fast” and “slow”: *l* = 0.4 and *l* = 1.0; “Transient:” and “sustained:”: *v* = 0.65 and *v* = 1.2. The resulting spike trains for the simulated cells, consisting of eight types, are shown in Fig. 1 together with each cell’s RF, the stimulus, linear response to the stimulus, and instantaneous spike rate.

##### RF parameter variation

To simulate cellular variability, each of the baseline RF parameters was varied by sampling from a Gaussian distribution centred around the baseline value, and constrained to positive values. The standard deviation was varied as a percentage proportion of the baseline value, using 5, 10, 15, 20, and 30% of each of the baseline parameters.

##### Imbalanced populations

To take into account that different RGC types are found in unequal numbers, imbalanced data sets were created. As representative examples, cell type frequencies were manipulated based on %-ON and %-fast, %-ON and %-transient, and %-fast and %-transient, in steps of 10%: %-ON from 30–70%, %-fast from 10–90%, and %-transient fixed at 50%; %-ON from 30–70%, %-transient from 10–90%, and %-fast fixed at 50%; %-transient from 30– 70%, %-fast from 10–90%, and %-ON fixed at 50%. This yielded 120 distinct data sets.

##### Recording noise

As perfect spike detection and sorting cannot be fully guaranteed in MEA recordings, the LNP model was augmented to include three sources of experimental noise:

1. Poisson process with a fixed rate throughout the stimulus time, either with a rate of *β* = 2*s^−^*1, or with a rate randomly chosen uniform between 5 Hz and 30 Hz. This simulates noise contamination of otherwise well isolated units.
2. Randomly removing 70% of the unit’s spikes across all trials, simulating false negatives during detection and/or spike sorting.
3. Merging two spike trains from different RFs, to simulate failed single unit isolation.

To generate datasets polluted with a known amount of noisy units, a random subset of the population was picked and replaced with one of the four noise models. The number of noisy units was evenly distributed between each of the four types. The percentage of noisy units as a percentage of the total population was varied from 10% to 90%, in increments of 10%, and a dataset was generated for each.

#### Experimental data

##### MEA recording

All experimental procedures were approved by the ethics committee at Newcastle University and carried out in accordance with the guidelines of the UK Home Office, under control of the Animals (Scientific Procedures) Act 1986. In this study, recordings from six C57bl/6 mice aged 59-101 days postnatal, housed under a 12 hour light-dark cycle, were used. All experimental procedures are described in detail in Hilgen et al. (21). Pan-retinal recordings were performed on the BioCam4096 platform with BioChips 4096S+ (3Brain GmbH, Lanquart, Switzerland), integrating 4096 square microelectrodes (21 by 21 μm, pitch 42 μm) on an active area of 2.67 by 2.67 mm. The platform records at a sampling rate of 7.1 kHz/electrode when using the full 4096 channel array and recordings were stored at 12 bits resolution per channel with a 8 kHz low-pass filter/0.8 kHz high-pass filter using 3Brain’s BrainWave software. Light stimuli were projected onto the retina as described previously (21) and attenuated using neutral density filters to high mesopic light levels (mean luminance 11 cd/m^2^).

##### Spike sorting

Spikes were detected and sorted using the algorithms described in (20, 29), using the HS2 software (https://github.com/mhhennig/HS2). Briefly, spikes were first detected as threshold crossings individually on each channel, and then merged into unique events based on spatial and temporal proximity. For each detected spike, a location was estimated based on the signal centre of mass. Spike sorting was performed by clustering all events using a feature vector consisting of the locations and the first two principal components of the largest waveform.

##### Analysis of light responses

The bias index (30) was computed from full field responses as

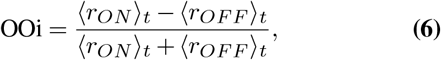

where 〈*r_ON_*〉_*t*_ 〈*roFF*〉 are the spike count during the bright and dark part of the stimulus, respectively.

DS was assessed by first recording the maximum spike count for each of 12 directions of a moving this by average across all stimuli and mean spike count of a neuron relative to its minimum spike rate for each of the moving bar stimuli:

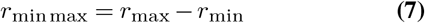

where *r*_max_ is the maximum and *r*_min_ is the minimum spike rate, calculated by counting the number of binned spikes, in this case using 200 ms wide bins. Taking the relative rate was required as it was found that the mean firing rate of the units drifted across the different moving bar stimuli, regardless of their actual peak response. The relative maximum rate was transformed into a vector representing the direction of the stimuli, and the first two normalised eigenvalues were used to compute the DSi:

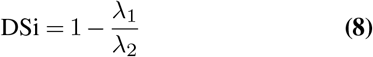

This value is bounded [0, 1], with a higher value signifying stronger DS.

The linear receptive field was computed as the spike triggered average (STA) stimulus, obtained during white noise stimulation. To increase the spatial resolution, the stimulus squares were shifted randomly with each presentation, as described by Hilgen et al. (20). A bivariate Gaussian was fitted to each STA at the time of the first peak, and its average width taken as the receptive field size.

##### Unit selection

To avoid false negatives during detection, a low detection threshold was used (threshold 10, see (29) for definition), which led to an excessive high number putative units. To isolate well-sorted neurons from this set, several heuristic criteria were applied: First, to be included, the eccentricity of the ellipse defined by a bi-variate Gaussian fit to the spatial spike locations of each unit was thresholded to *<* 0.85, and the average of the two axes thresholded to below <17% of the channel separation (7.14μm). This removed units with poorly localized, and therefore potentially poorly sorted units. Units recorded from the two outer channel rows were removed as this impaired spike sorting Next, only units with at least 10 spike in eah trial of the chirp stimulus were retained. Here we used six retinas, with a final count of 1026, 1849, 1234, 634, 1131 and 575 units.

### Results

Our procedure to obtain functional RGC clusters consists of two steps. First a spike distance matrix is computed for all single unit pairs in the recording, which is then clustered using agglomerative clustering. This requires a distance measure with the metric properties of non-negativity, zero distance for identity, symmetry and subadditivity (the sum of two distances has to be greater than or equal than the individual distances). These conditions are fulfilled by a number of metrics (22, 24, 31), of which we evaluated the non-parametric ISI and SPIKE distance measures. Then hierarchical clustering is used to construct a dendrogram based on the distance matrix. In the following, we will first validate this method with synthetic data, and then show its application to a high density MEA recording from the mouse retina.

### Evaluation with synthetic data

We first evaluate the method with a synthetic ground-truth data set, which contains eight different RGC types (see Methods). Spike trains were generated using LNP models, and the spike distance matrix was computed, followed by hierarchical agglomerative clustering using Ward’s linkage. This procedure yields a dendrogram, which was cut such that eight flat clusters were obtained. We note that the last step is not feasible for recorded spike trains, where the number of RGC types is not known. We address this issue below when analysing an experimental data set.

First we combine all 140 datasets created using the LNP model. In addition to the spike distance-based methods, the ISI and SPIKE distances, we also used the raw PSTH, and the PCA and sparse PCA of the PSTH for comparison (see Baden et al. for an application of sparse PCA). To assess the quality of the clustering, the median score of the following four measures was used: the adjusted Rand index, which summarises the fraction of correct clustering choices based on true/false negatives/positives (32); the adjusted mutual information, which quantifies information gain over random clustering (33); the Fowlkes-Mallows score, the geometric mean of precision and recall; and the completeness score measuring if all true cluster members are in the same cluster. Across almost all conditions, the scores of each individual quality index were very similar. The distribution of the median scores across all conditions for each method is shown in Fig. 2.

**Fig. 2.**
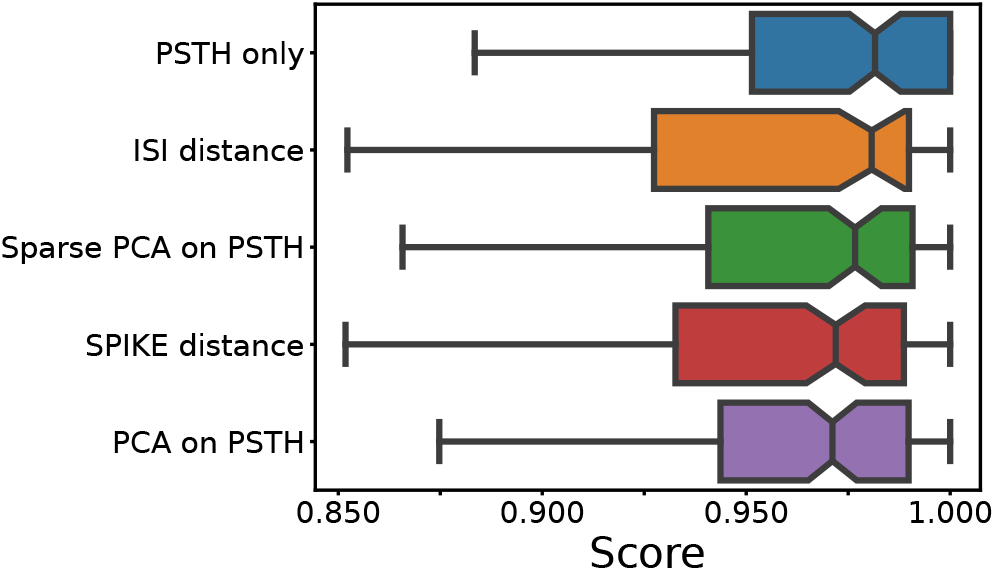
Median score across all conditions. For each of the five feature sets, four external clustering quality scores (adjusted Rand index, adjusted mutual information score, V-measure, and the Fowlkes-Mallows score) were computed for each dataset. The median of the four scores is calculated for each dataset. Finally, the medians for the various datasets (140) are presented in the form of a box plot, one for each method. A higher score implies a better clustering result. Each coloured box plot shows the median and interquartile range (IQR). The error bars extend 0.8 times past the IQR in each direction. The methods are ordered from the highest median value (of median scores) to the lowest, from top to bottom.

This comparison shows that the different feature sets yield a similar performance. However, it should be noted that the methods using PSTHs, either alone or through a form of dimensionality reduction, require selecting and optimising the PSTH bin size. Secondly, PCA requires selecting the dimensionality, and sparse PCA has an additional L1 penalty parameter. An exploration of the the ground truth data showed that all these parameter strongly affect clustering quality. For the results shown in Fig. 2, the parameters optimised through an extensive grid search.

For the PSTH-based clustering, a bin size of 200 ms resulted in the best overall performance, which also proved to be the best bin size for PCA and sparse PCA. Note that this large bin size was likely required due to the high variability introduced by the Poisson spike generation process. In real data, more precise spiking will require a smaller bin size. For PCA, using eight PCs worked best, whereas for sparse PCA, a penalty of 50 with 12 components provided the best scores.

In contrast, the ISI and SPIKE distance based clustering required no such parameter search. The fact that both methods have comparable or better performance demonstrates that they are suitable for the clustering of RGC types based on their physiological response. This is particularly important in the context of real data, where parameter optimisation can only be done heuristically.

### Synthetic data with biological variability

Comparing the methods as a function of cell number and RF variation, we find that the performance degraded rapidly as the RF variation increased beyond 15% (Fig. 3). In particular, the spike distance based measures performed slightly worse than the others as RF variation increased. In all cases, the worst performance is observed when the RF variation is at 30%, and the total number of units is high (800 units). Each of the methods performs consistently with regard to its relative overall performance (the order shown in Fig. 2). This trend is consistent across all conditions of the synthetic dataset: permitting for minor local variations, there was no single direction in which any of the methods described performed characteristically better or worse.

**Fig. 3.**
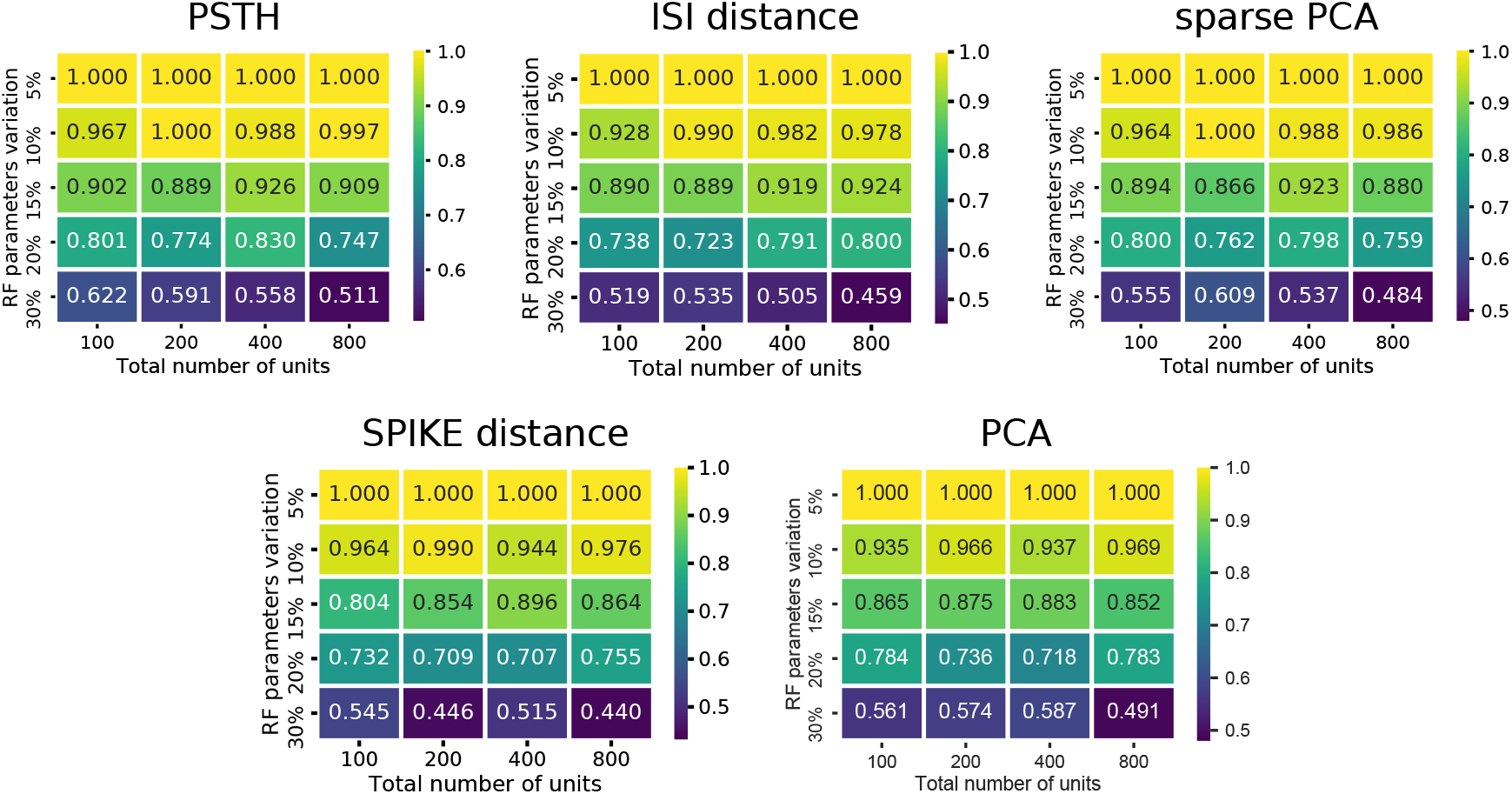
Cluster quality measures as a function of RF variation and total number of units. For each of the five feature sets, the median of four clustering quality scores (adjusted Rand index, adjusted mutual information score, V-measure, and the Fowlkes-Mallows score) are shown for each dataset (140 in total).

### Synthetic data with experimental noise

To evaluate the effect of experimental noise, we systematically varied the proportion of noisy units (see Methods). For the three methods requiring parameter selection, the values that gave the best results on the non-noisy synthetic data were used. Here, only the labels of the ‘good’, noise-free, units were compared between the clustering results and the original labels. Therefore, the comparison does not assess the methods for their ability to separate the ‘bad’ noisy units from the ‘good’ ones. Instead, the emphasis is purely on being able to separate the different types of RGCs from each other, even if noisy units belong to these clusters. The rationale for this choice is that if an RGC recording has been properly pre-processed and filtered, the proportion of bad units should be relatively low, and, therefore, the mean response of an RGC cluster should not be affected. Moreover, the noise model introduces new ‘types’ of cells, hence constraining the results to eight flat clusters could force erroneous clustering results. Therefore the results were evaluated against a range of numbers of flat clusters, representing different cutoff points within the hierarchy.

The median score across methods obtained for a clustering result of eight flat clusters show a significant degradation in reconstruction quality as the number of noisy units increases (Fig. 4). The ISI and SPIKE distance measures in particular appear to deteriorate immediately upon the introduction of bad units, despite a comparable baseline. Closer inspection of the scores shows that the completeness score in particular reveals an interesting result. This score places emphasis on classes being grouped together, and will result in a score of 1.0 even if all classes are part of a single cluster. The high completeness scores for the spike distance measures, in particular the SPIKE distance, for a high fraction of noisy units shows they correctly group RGC types with remarkable accuracy (Fig. 4). This indicates that the SPIKE distance clustering separates the noisy units into their own clusters, and, as a result of the cutoff point in the hierarchy, a flat clustering of 8 groups is insufficient to capture all types. This leads to lower scores on the other measures, but would allow recovering a correct clustering when more flat clusters are allowed.

This is indeed the case when creating 16 flat clusters (Fig. 4). While the median score of all the external clustering quality measures improves, the median quality score for the baseline case (0% bad units) is now, as expected, lower for all methods. However, the ISI and SPIKE distance methods improve dramatically for low to medium levels of noise (10–60%), with SPIKE distance surpassing all others over this range. Yet, both spike distance methods continue to fare poorly when the population is dominated by noisy units (70–90%). The completeness scores agree with these observations, but they go further by demonstrating that the reason for the poor performance in the high noise range does not affect its ability to continue to correctly group RGC types in highly contaminated data.

**Fig. 4.**
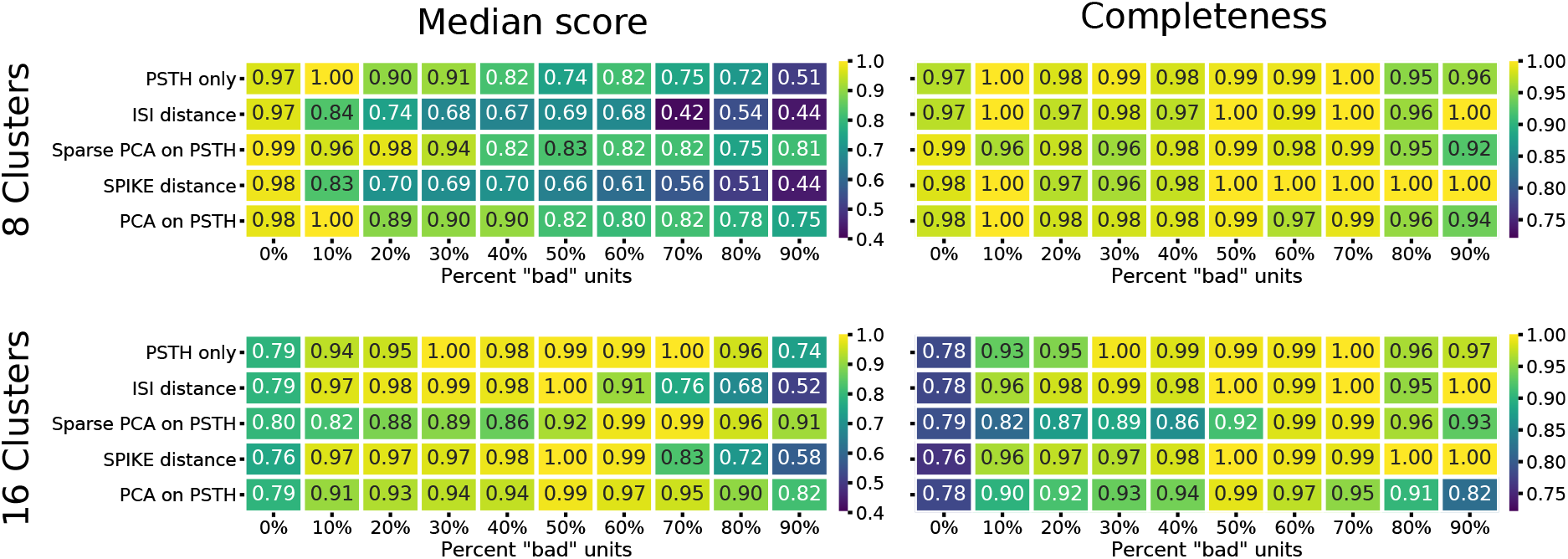
Clustering quality scores on noisy data. For each of the five methods, the median of four external clustering quality scores (adjusted Rand index, adjusted mutual information score, V-measure, and the Fowlkes-Mallows score) are shown in the left panels. Right panels show the completeness score. The top row show the results for 8, and the bottom row for 16 flat clusters.

In sum, these results show that spike distance measures can be used to classify RGC responses. In comparison with other methods, the procedure does not depend on any parameters, which would have to be chosen heuristically and can significantly affect the clustering results. In addition, the SPIKE distance in particular is robust to experimental noise, which is hard to avoid in large scale, high-density recordings where manual data curation is infeasible. High completeness scores indicate that clusters are not wrongly mixed in this case, but instead noisy units form separate clusters that can be identified manually.

### Clustering experimental data

Next we used the spike distance measures to cluster RGC types from one mouse retina high-density MEA recording. Following quality control (see Methods), a set of 1026 well isolated RGCs was included in this analysis. Light-evoked activity used to compute the distance matrices was obtained during stimulation with either full field flashes, or a stimulus that included a full field contrast step, green and blue full field flashes and a chirp sequence (see Methods).

Hierarchical agglomerative clustering constructs a dendrogram, starting with each single unit as its own cluster, and iteratively merging units and resulting clusters with a minimum variance constraint. As we have shown for synthetic data, the spike distance measures are very robust to variability and noise. However, to ensure noisy units are indeed placed into their own clusters while preventing over-clustering, an appropriate dendrogram cut-off point has to be determined. We evaluated two common metrics: The first was the gap statistic, which compares the compactness of each cluster to a random surrogate (34). Using the shuffled distance matrix as surrogate, this metric indicates the data contains between 17 and 18 clusters, depending on the stimulus and metric (Fig. 5A). These numbers are similar to those reported by Farrow and Masland (15), but fall short of the higher numbers given by Baden et al. (17). However, we found that the gap statistic generally favours larger clusters in data with higher variability, as small clusters naturally yield squared distances closer to the random surrogate. Hence the gap statistic may be a measure too conservative to reliably assess data with potentially highly imbalanced clusters.

**Fig. 5.**
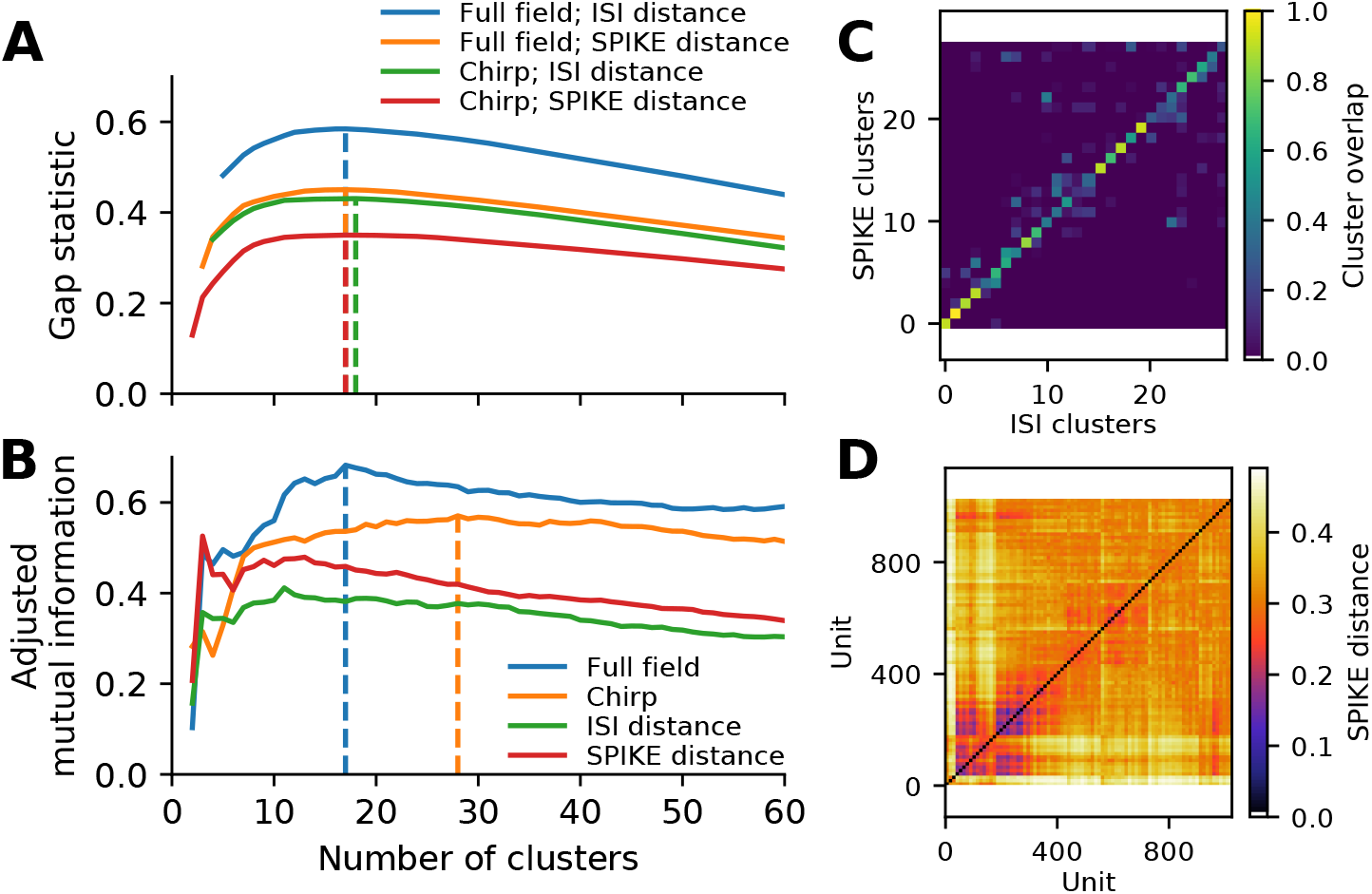
Cluster metrics for a recording from 1026 retinal ganglion cells. **A** The Gap statistic for the two spike distance metrics, computed for the full field and chirp stimulus, as a function of the number of flat clusters. Vertical lines indicate the peak of each curve. **B** Paired comparison of different clustering outcomes, quantified using adjusted mutual information. Two pairings were performed: between the ISI and SPIKE distance metrics for full field (blue) and chirp (orange), and between the two stimuli (full field and chirp) for either ISI (green) or SPIKE (red) distance. Vertical lines indicate the peak values for the comparison between spike distances. **C** Overlap between clusters obtained with the ISI and SPIKE metric, for 31 flat clusters. **D** The SPIKE distance matrix for all units, ordered by linkage.

To see whether the data may contain more valid clusters, we turned to a consensus method. To this end, we compare the results obtained with the ISI and SPIKE distance for the two stimuli, using adjusted mutual information (33). This peaked at 17 clusters for the full field stimulus, and 28 clusters for the chirp stimulus (Fig. 5B). Investigating this metric and the corresponding confusion matrices (see Fig. 5C for the case of 28 clusters), it is evident that consensus is high for solutions with 15 or more clusters for both stimuli. In addition, the SPIKE distance consistently scored higher for a comparison between full field and chirp stimulus. Therefore, in the following we chose the SPIKE distance, using the peak conensus value of 28 clusters.

To get a first impression of the clustering result, we used t-distributed Stochastic Neighbor Embedding (t-SNE) to create a two-dimensional, non-linear embedding of the chirp PSTHs for visualisation (Fig. 6). In this plot, each dot represents a single cell, and spatial proximity indicates high similarity. Note however that the overall spatial arrangement is arbitrary and that distance does not imply similarity.

**Fig. 6.**
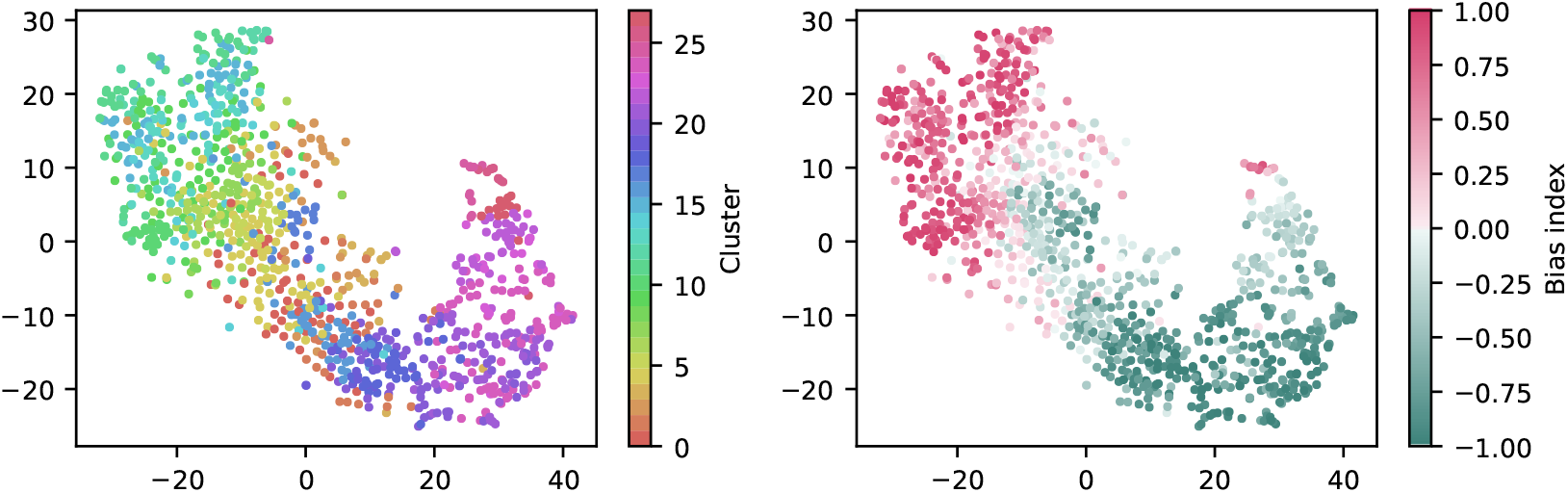
Visualisation of the clustering with t-SNE embeddings of the chirp responses. A t-SNE embedding was generated from chirp PSTHs (bin size 50ms; perplexity 30). Each unit is coloured either by cluster membership (left) or by bias index (right). Note that similar colors in the cluster plot are perceptually hard to distinguish, colours were chosen such that a similar hue indicates proximity in the cluster dendrogram. The colour coding is the same as in Fig. 7.

Colouring each dot by cluster membership reveals that the area covered by each cluster is limited (Fig. 6, left). This suggests the clustering captures relevant structure in the data. Moreover, colouring the same plot by the bias index shows that it is a major contributor to similarity (Fig. 6, right), a feature well captured by the clustering (note that this information was not included explicitly in the clustering process). It is noteworthy that the same analysis for the full field response yields a less structured picture, indicating that the richer chirp stimulus is more informative about specific RGC types (not illustrated).

Finally, a summary of the resulting 28 clusters is shown in Fig. 7. The dendrogram, which is coloured to indicate the average bias index on each branch (green = Off; red = On; grey = no preference) is organised into three main branches. The first contains many On cells and a sub-branch with mixed preference. The second branch contains exclusively Off cells. The third branch has again mixed preference, with an average bias index close to zero. Common to RGCs in the last branch appears to be transient On or Off responses with a stronger sustained component.

**Fig. 7.**
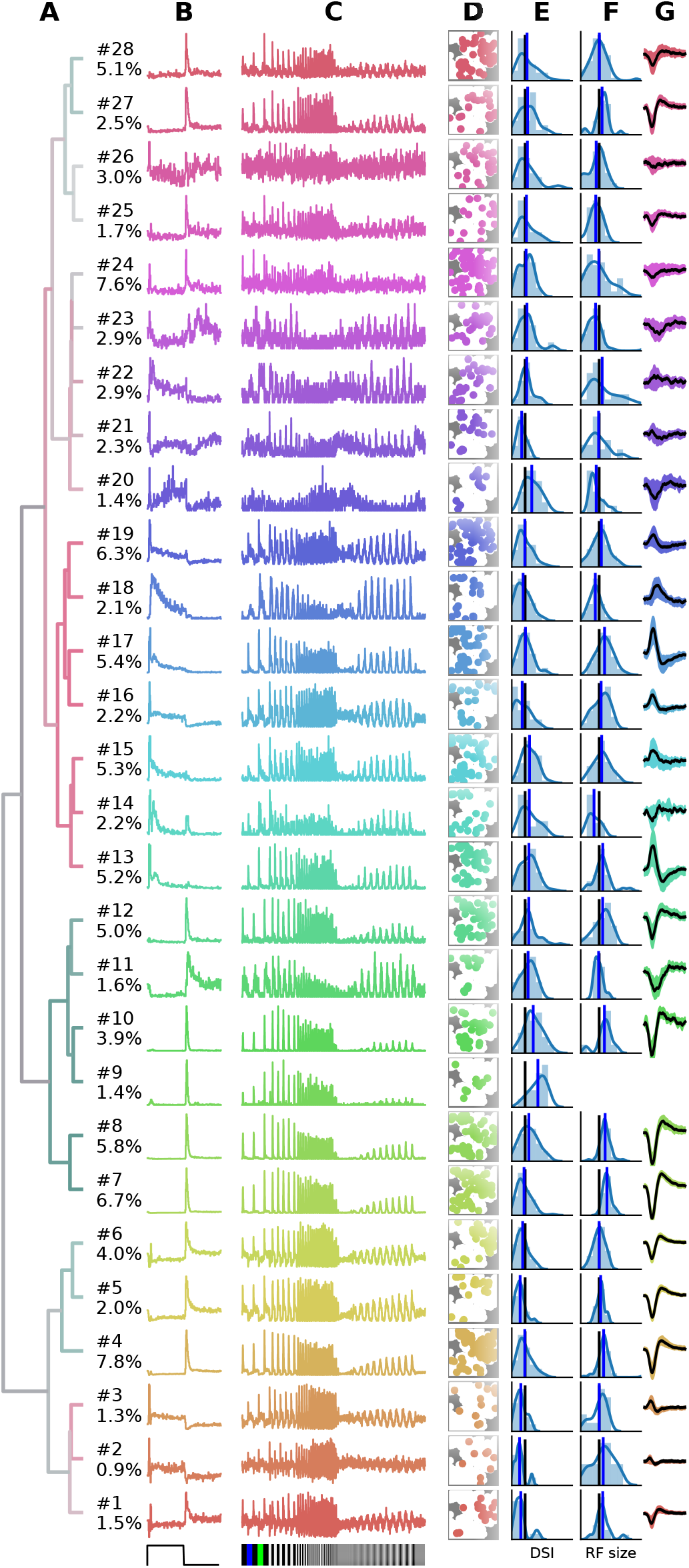
Summary of the RGC types found in a recording from 1026 retinal ganglion cells (RGCs). **A**, Dendrogram, with lines coloured according to the average bias index of all units with lower distance (green=-1/Off, red=+1/On, grey=Zero). Percentages at the leafs give the relative abundance of each RGC type. **B**-**G**, Each row shows properties of one cluster: **B**, full field PSTH (2 sec bright, 2 sec dark); **C**, chirp PSTH; **D**, spatial distribution of each cluster, recorded area is indicated in white; **E**, histogram of direction selectivity index (DSi); **F**, histogram of receptive field sizes; **G**, temporal spike triggered average (STA) obtained at the pixel with the largest STA magnitude, shaded area indicates one standard deviation around the mean (black, where missing no receptive field could be fit). In E and F, the black vertical lines are retina averages and blue lines the cluster averages.

While it is not the aim of this work to exhaustively characterise the RGC types found in this experiment, in the following some interesting cases are discussed. First, the PSTHs exhibit clear differences in response kinetics temporal and contrast sensitivity between clusters. In some cases, these differences are rather subtle, for instance clusters 7 and 8 appear superficially similar, and both could be classified as the highly abundant Off alpha cells with transient responses and large RFs. Yet cluster 7 has a slightly higher sustained response and a higher frequency and contrast sensitivity. Cluster 8 may therefore correspond to the “OFF mini alpha transient”, while 7 may be the “Off alpha transient” type as described by Baden et al. (17). To fully confirm their identify, a more detailed analysis would be required.

Fig. 7 also shows spatial distribution, DS, RF size estimated from STAs, and the temporal STA profile for each cluster. Direction selectivity and RF size generally show broad distributions, but with consistent differences between types. For instance, the characteristics cells with larger than average RFs match types with known large RFs, such as Off-alpha like cells (clusters 7, 8, 10 and 12), and a putative On alpha type in cluster 17 (13). On the other hand, several clusters have very weak (2, 14, 15, 21-26) or even no well defined (cluster 9) STA, adding variability in the RF fits.

Direction selectivity was measured with a single bar moving in 12 different directions, a paradigm we found to be relatively susceptible to noise. It is therefore unclear if the high variability in many clusters is due to experimental noise, or due to incomplete separation of DS types. Two observations however stand out. First, all cells grouped together into the third mixed supercluster (clusters 1-6) have a very low DS, lower than any other cluster. Second, cluster 9 has a very strong DS. None of the RGCs in this cluster had a measurable STA, suggesting it may correspond to the “ON–OFF DS 2” type (17).

Finally, cluster 26 does not appear to have a discernible light response. Closer inspection revealed that it consists of noisy units with highly inconsistent firing patterns. Hence these cluster are not likely to contains reliable RGC activity, but one or several poorly detected neurons. It is encouraging that, as for the synthetic data, these noisy units are sorted into separate clusters, which allows easy removal from further analysis.

### Heterogeneity between retinas

To assess the consistency of RGC classification across recordings from different retinas, we performed pairwise comparisons between six retinas. These had 1026, 1849, 1234, 634, 1131 and 575 units, respectively (retina 1 is the same as shown in Fig. 7). Both the gap statistics and the adjusted mutual information show a similar dependence on the cluster number as shown above for one retina, but vary in their peak values, indicating differences in overall separability in each preparation (not illustrated). Each retina was clustered separately, and a PCA model was fit using the combined, peak-normalised chirp PSTHs from all retinas. Matching clusters between each pair of retinas were found based on the smallest cosine distance between the eight first components. The cosine distance, which is best suited to compute distances in a vector space, ranges from zero to one, where zero indicates identity and two opposing vectors. As expected, averaged distances decrease as the number of clusters is increased, but remains high even for 70 clusters (Fig. 8A). In addition, we found considerable variability in distances between retinas, indicating that responses from the same RGC type can be variable across retinas.

**Fig. 8.**
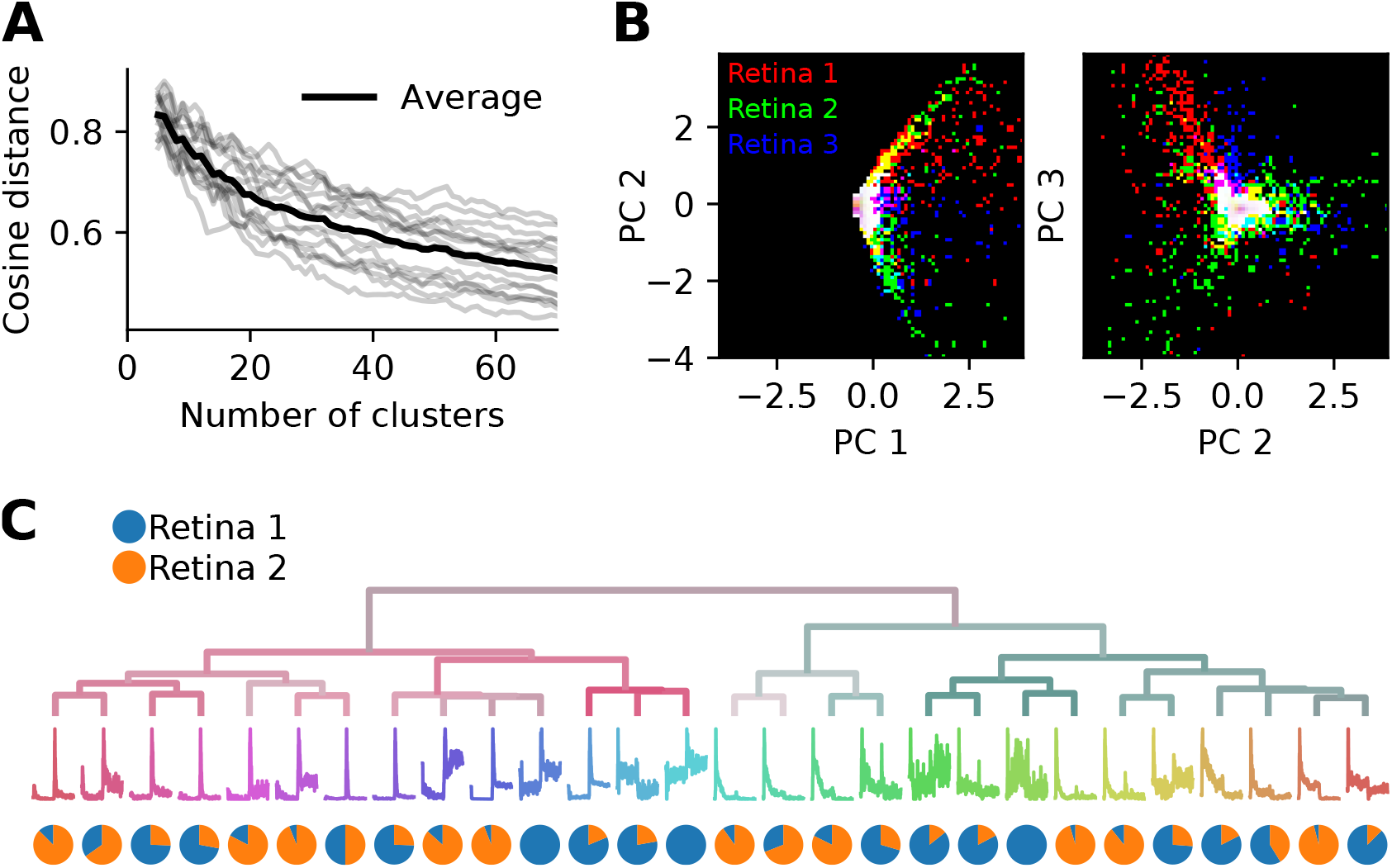
Systematic differences in RGC activity between individual retinas prevents data pooling. **A**, Grey lines show, for each pair of six retinas, the average cosine distances between the top eight PCA projections of the best matching chirp PSTHs (explaining 78% variance), as a function of the number of clusters generated. The black line is the average across all data sets. Individual retinas show high variability even for a large number of clusters, indicating considerable differences between preparations. **B**, The first and second (left) and second and third PCA projections plotted against each other for three retinas (indicated by colour). Each retina occupies a different region in PCA space.**C**, Dendrogram obtained by clustering a data set generated from randomly selecting 500 neurons from two retinas. The pie charts show the relative proportion of neurons from each retina in each cluster. Most clusters exhibit a strong bias towards one of the two retinas.

This effect can also be seen when visualizing the principal components directly for each retina. Fig. 8B illustrates this for the first three components for three retinas. While some regions in PCA space are shared, each retina occupies distinct areas. As a result, SPIKE distance-based clustering of spike trains pooled from different retinas yields results that primarily separate units by retina, and not by functional type. This is illustrated in Fig. 8C, where a mixed data set was generated by randomly sampling 500 neurons from retinas 1 and Many of the resulting clusters contain neurons from only one of the two retinas.

The variability between preparations complicates finding matching clusters across all retinas, which we attempted using a greedy search procedure. However, for pairs of retinas this was feasible to some extent, as distances were similarly biased for the corresponding cluster pairs. To find matching pairs, kernel PCA using radial basis functions was fit with 20 components to peak-normalised chirp PSTHs, and cosine distances were computed between all neurons for each pair of clusters and averaged. This procedure was carried out using two retinas, clustered into 34 types each instead of the 28 clusters used above. We chose this number as the synthetic data experiments showed that noisy neurons are assigned their own clusters, while the main types are still preserved. Hence over-clustering is not expected to happen unless this number is increased drastically.

When pairing these clusters by shortest average distance, we found 27 clusters with well matched bias indices, and with correlated direction selectivity (Fig. 9A). Well matched clusters tended to have small distances, while they were larger for mismatched clusters. The distances increased sharply after 24 clusters (Fig. 9B), indicating that at least 10 clusters could not be matched well. Fig. 9C illustrates a direct comparison of these 24 clusters.

**Fig. 9.**
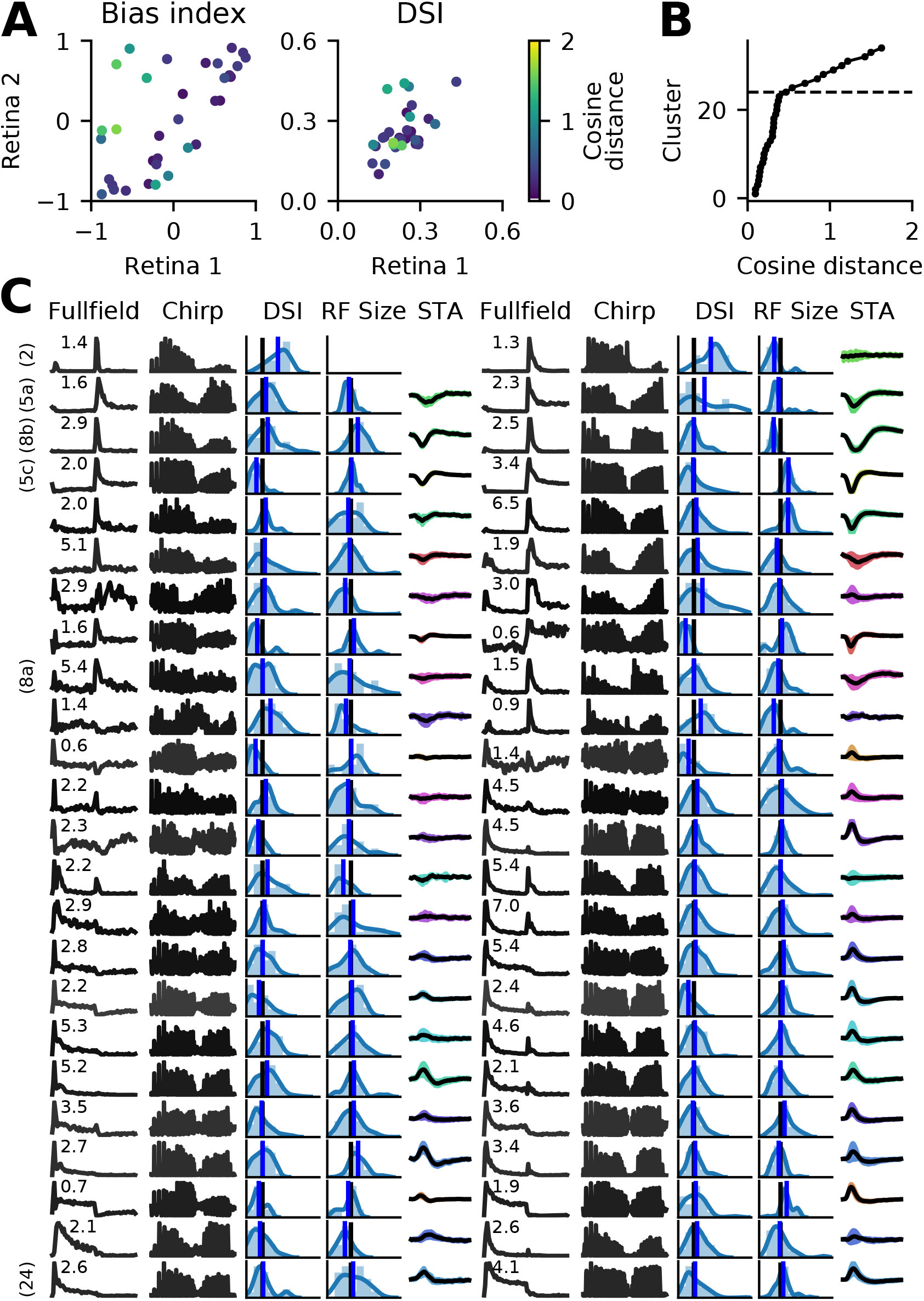
Matching RGC types in two retinas. Spike trains from two retinas (retinas 1 and 2 in Fig. 8) were independently clustered into 34 types, and matching pairs found using the cosine distance between vectors of length 20, obtained with RBF kernel PCA. **A**, A comparison of average bias indices (left) and direction selectivity (right) for each pair of RGC types. Colour indicates the distance measure for each pair. **B**, Cumulative histogram of average pair distances. Beyond 24 clusters, a sharp increase in distance is observed. **C**, Main features of the 24 most similar RGC types for both retinas. Data is presented as in Fig. 7, with full field and chirp PSTHs, histograms of direction selectivity index (DSi) and receptive field size (black vertical lines are retina averages, blue line is the cluster average), and the STA (shaded area: 1 standard deviation; missing STA could not be fit). Numbers above the full field PSTHs are the percentage abundance of each RGC type. PSTHs are coloured by cosine distance, with black indicating zero and grey higher values. On the left, clearly identified RGC types according to the taxonomy by Baden et al. (17) are given.

For six of these clusters, we could identify corresponding counterparts in the taxonomy developed by Baden et al. (17): Off DS type without clear STA (group number 2 in Baden et al. (17)), Off alpha sustained (5a and 5c), Off alpha transient (8a, 8b) and On alpha (24) (annotated on the left in Fig. 9C). However, a direct comparison to other classification studies is difficult because of the nature of the different acquisition conditions. To illustrate the variability within each retina, and between retinas, PSTHs of all neurons for each of these annotated clusters are shown in Fig. 10. While overall consistent, there are clear differences in firing rates between retinas, and for the same type within the same retina. The origin of this variability is unclear, possible factors include experimental limitations (coupling of the retina to the MEA, variability in spike detection and sorting), differences between the retinal eccentricities and recording locations, and biological neuronal heterogeneity.

**Fig. 10.**
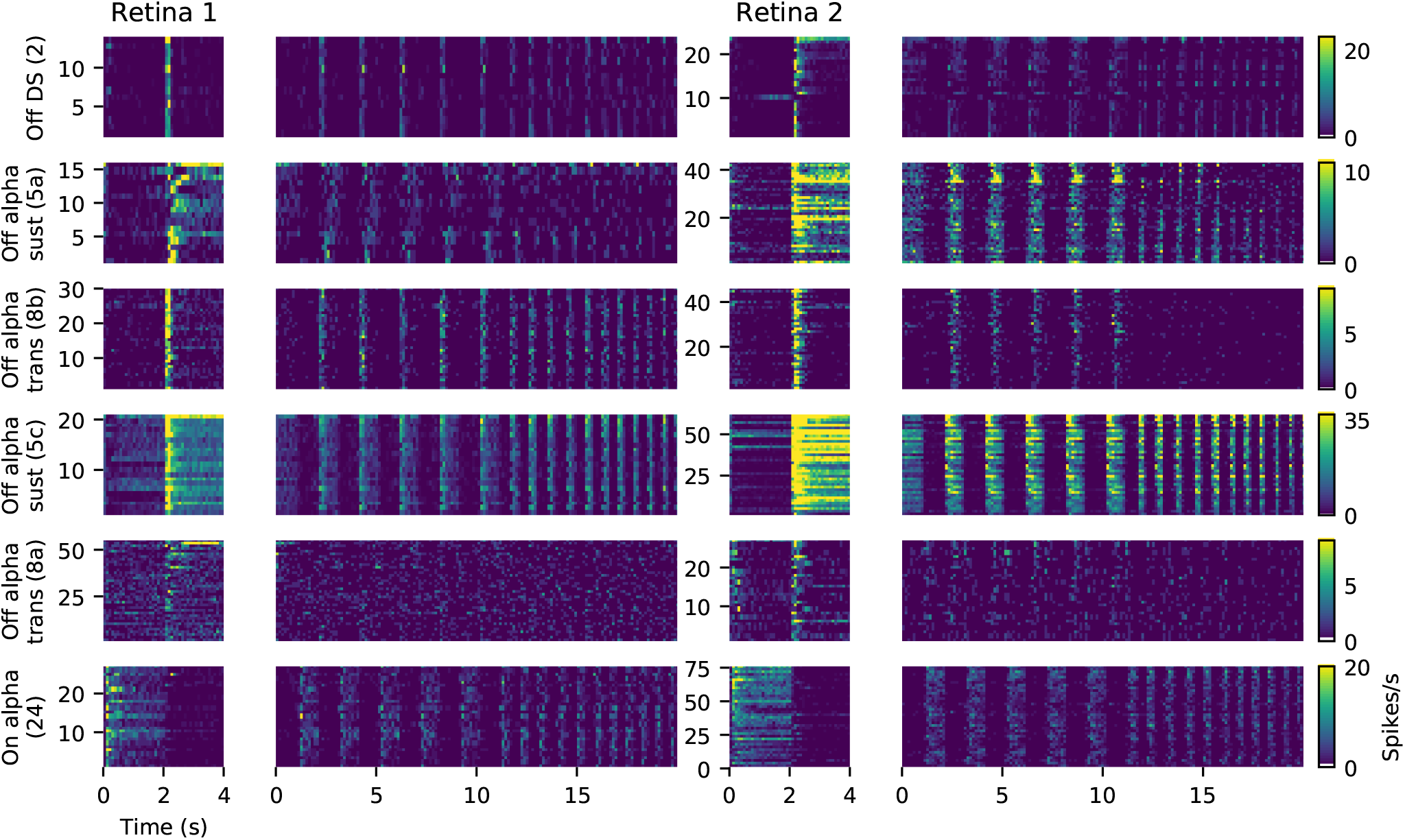
Examples of matched cell types in two retinas. Each panel shows the PSTHs during full field stimulation (small, left) and the frequency-modulated part of the chirp stimulus (right), for all neurons clustered together as the same type. Data is the same as shown in Fig. 9. The colour gradient indicating firing rates is peak-scaled to the largest response across all neurons in retina 1 for the chirp stimulus, separately for each type.

## Discussion

This work introduces a method for classification of RGC responses based solely on spatially uniform light stimulation, using spike train distances as a metric for hierarchical clustering. The analysis of synthetically generated ground truth data shows that this method performs well. For noisy data, our method outperforms other approaches, which typically perform clustering in a low-dimensional Euclidean feature space obtained through dimensionality reduction (9, 15, 30, 35, 36). Most importantly however, our method is entirely non-parametric, thus does not require searching for optimal hyperparameters, such as the dimensionality of the feature representation (17), or the definition or estimation of salient response properties.

Our approach extends the idea by Zeck and Masland (16) to use salient spike train features directly for clustering. This approach requires all neurons to be stimulated simultaneously with the same stimulus, to ensure the spike distance measure only captures physiological variations. This is a very effective way to probe many RGCs simultaneously with high density MEAs, since a small number of trials of a short, but sufficiently rich stimulus are sufficient. While only modulating contrast and temporal aspects of the RFs, the chirp stimulus appears well suited as it allows to rapidly probe a range of relevant response properties. Some clusters showed distinct receptive field sizes, high direction selectivity or no clearly defined STA, indicating that their responses to this homogeneous stimulus are sufficiently distinct to isolate sub-types also with non-optimal stimuli (16).

The output of our method is a dendrogram indicating similarity between units and groups of units. To obtain a flat clustering into distinct RGC types, an appropriate cut-off point has to be determined. This can be done either by computing a consensus score for two or more distance measures, or by manual inspection. Consistent with our simulations, we found that clusters with noisy, unreliable activity were split first when the number of clusters was increased, while well-identified RGC types remained in the same clusters. This allows the identification of frequent types, such as, for instance, Off-alpha cells or On mono- and bi-phasic RGCs. Noisy, invalid units, on the other hand, can be removed by first clustering a full, contaminated data set, then removing clusters with inconsistent responses, followed by re-clustering.

A further method to assess the number of RGC types is consensus clustering of recordings from different retinas. However, perhaps unsurprisingly, we found significant variability between retinas when comparing different recordings. This prevents the joint clustering of pooled data, as spike distance measures are highly sensitive to these systematic differences. As a result, the clusters obtained for pooled data frequently group neurons from the same retina. An alternative approach we explored here is to cluster the recordings individually, followed by computing pairwise distances between all recordings.

As pointed out before, light responses alone are certainly insufficient to conclusively distinguish between individual RGC types (17). A very reliable criterion is neural morphology, in particular dendritic stratification in the inner plexiform layer. While this is impossible to obtain for all neurons recorded with large scale MEAs, the registration of sparsely labelled neurons with spike sorted data is feasible (20), a method that could be used to augment the clustering obtained with our method. A further criterion to assess the validity of clustered types is regular spatial tiling of each type (35, 37), although there are exceptions to this rule (38). Indeed our analysis suggests some cell types may be preferentially found in certain regions of the retina (cf. Fig. 7D), a finding that however has to be substantiated by comparing recordings from the same area in multiple retinas.

Taken together, here we show that spike train distances can be successfully used to classify RGCs in the mouse retina. We expect this approach can be used in other sensory systems where an appropriate homogeneous stimulus can be delivered. A potential shortcoming is that cell types with a continuous variation of selectivity, such as auditory neurons with different frequency tuning, will be wrongly placed into different groups. We also expect our method to be suitable for analysis of spike trains inferred from calcium imaging recordings. The analysis of synthetic data shows that the method is robust even for highly variable Poisson spike trains, as long as key response features are well preserved. As also pointed out by Zeck and Masland (16), transient, well-timed responses are particularly important for spike-based measures, which we expect can be inferred well from clean imaging data. On the other hand, given the results discussed above, it seems unlikely our method will perform well for data pooled across multiple preparations.

## ACKNOWLEDGEMENTS

Funding was provided by the Engineering and Physical Sciences Research Council grant EP/L027208/1 (MHH), and the Leverhulme Trust grant RPG-2016-315 (ES).

## Bibliography

1. Botond Roska and Frank Werblin. Vertical interactions across ten parallel, stacked representations in the mammalian retina. Nature, 410(6828):583, 2001.

2. H Wässle, L Peichl, and BB Boycott. Dendritic territories of cat retinal ganglion cells. Nature, 292(5821):344, 1981.

3. R. L. Rockhill, T. Euler, and R. H. Masland. Spatial order within but not between types of retinal neurons. Proceedings of the National Academy of Sciences, 97(5):2303–2307, 2000. ISSN 0027-8424. doi: 10.1073/pnas.030413497.

4. Wenzhi Sun, Ning Li, and Shigang He. Large-scale morphological survey of mouse retinal ganglion cells. Journal of Comparative Neurology, 451(2):115–126, 2002.

5. Tudor Constantin Badea and Jeremy Nathans. Quantitative analysis of neuronal morphologies in the mouse retina visualized by using a genetically directed reporter. Journal of Comparative Neurology, 480(4):331–351, 2004.

6. Jee-Hyun Kong, Daniel R Fish, Rebecca L Rockhill, and Richard H Masland. Diversity of ganglion cells in the mouse retina: unsupervised morphological classification and its limits. Journal of Comparative Neurology, 489(3):293–310, 2005.

7. Béla Völgyi, Samir Chheda, and Stewart A Bloomfield. Tracer coupling patterns of the ganglion cell subtypes in the mouse retina. Journal of Comparative Neurology, 512(5): 664–687, 2009.

8. Richard H. Masland. The Neuronal Organization of the Retina. Neuron, 76(2):266–280, oct 2012. doi: 10.1016/j.neuron.2012.10.002.

9. Ian L Jones, Thomas L Russell, Karl Farrow, Michele Fiscella, Felix Franke, Jan Müller, David Jäckel, and Andreas Hierlemann. A method for electrophysiological characterization of hamster retinal ganglion cells using a high-density cmos microelectrode array. Frontiers in neuroscience, 9:360, 2015.

10. Moritz Helmstaedter, Kevin L Briggman, Srinivas C Turaga, Viren Jain, H Sebastian Seung, and Winfried Denk. Connectomic reconstruction of the inner plexiform layer in the mouse retina. Nature, 500(7461):168, 2013.

11. Evan Z Macosko, Anindita Basu, Rahul Satija, James Nemesh, Karthik Shekhar, Melissa Goldman, Itay Tirosh, Allison R Bialas, Nolan Kamitaki, Emily M Martersteck, et al. Highly parallel genome-wide expression profiling of individual cells using nanoliter droplets. Cell, 161(5):1202–1214, 2015.

12. Joshua R. Sanes and Richard H. Masland. The Types of Retinal Ganglion Cells: Current Status and Implications for Neuronal Classification. Annual Review of Neuroscience, 38: 221–246, 2015. ISSN 0147-006X. doi: 10.1146/annurev-neuro-071714-034120.

13. Brenna Krieger, Mu Qiao, David L Rousso, Joshua R Sanes, and Markus Meister. Four alpha ganglion cell types in mouse retina: Function, structure, and molecular signatures. PloS one, 12(7):e0180091, 2017.

14. Bruce A Rheaume, Amyeo Jereen, Mohan Bolisetty, Muhammad S Sajid, Yue Yang, Kathleen Renna, Lili Sun, Paul Robson, and Ephraim F Trakhtenberg. Single cell transcriptome profiling of retinal ganglion cells identifies cellular subtypes. Nature communications, 9(1): 2759, 2018.

15. K. Farrow and R. H. Masland. Physiological clustering of visual channels in the mouse retina. Journal of Neurophysiology, 105(4):1516–1530, 2011. ISSN 0022-3077. doi: 10. 1152/jn.00331.2010.

16. Günther M. Zeck and Richard H. Masland. Spike train signatures of retinal ganglion cell types. European Journal of Neuroscience, 26(2):367–380, 2007. ISSN 0953816X. doi: 10.1111/j.1460-9568.2007.05670.x.

17. Tom Baden, Philipp Berens, Katrin Franke, Miroslav Román Rosón, Matthias Bethge, and Thomas Euler. The functional diversity of retinal ganglion cells in the mouse. Nature, 529 (7586):345–350, jan 2016. ISSN 0028-0836. doi: 10.1038/nature16468.

18. Alessandro Maccione, Matthias Helge Hennig, Mauro Gandolfo, Oliver Muthmann, James van Coppenhagen, Stephen J. Eglen, Luca Berdondini, and Evelyne Sernagor. Following the ontogeny of retinal waves: pan-retinal recordings of population dynamics in the neonatal mouse. Journal of Physiology, 592(7):1545–1563, 2014. ISSN 14697793. doi: 10.1113/ jphysiol.2013.262840.

19. Geoffrey Portelli, John M Barrett, Gerrit Hilgen, Timothée Masquelier, Alessandro Maccione, Stefano Di Marco, Luca Berdondini, Pierre Kornprobst, and Evelyne Sernagor. Rank order coding: a retinal information decoding strategy revealed by large-scale multielectrode array retinal recordings. Eneuro, pages ENEURO–0134, 2016.

20. Gerrit Hilgen, Sahar Pirmoradian, Daniela Pamplona, Pierre Kornprobst, Bruno Cessac, Matthias Helge Hennig, and Evelyne Sernagor. Pan-retinal characterisation of Light Responses from Ganglion Cells in the Developing Mouse Retina. Scientific Reports, 7(42330): 42330, feb 2017. ISSN 2045-2322. doi: 10.1038/srep42330.

21. Gerrit Hilgen, Martino Sorbaro, Francesca Cella Zanacchi, Evelyne Sernagor, Matthias Helge Hennig, Sahar Pirmoradian, Jens-Oliver Muthmann, Ibolya Edit Kepiro, Simona Ullo, Cesar Juarez Ramirez, Albert Puente Encinas, Alessandro Maccione, Luca Berdondini, Vittorio Murino, and Diego Sona. Unsupervised Spike Sorting for Large-Scale, High-Density Multielectrode Arrays. Cell Reports, 18(10):2521–2532, mar 2017. ISSN 22111247. doi: 10.1016/j.celrep.2017.02.038.

22. Thomas Kreuz, Julie S. Haas, Alice Morelli, Henry D.I. Abarbanel, and Antonio Politi. Measuring spike train synchrony. Journal of Neuroscience Methods, 165(1):151–161, 2007. ISSN 01650270. doi: 10.1016/j.jneumeth.2007.05.031.

23. Thomas Kreuz, Daniel Chicharro, Martin Greschner, and Ralph G. Andrzejak. Time-resolved and time-scale adaptive measures of spike train synchrony. Journal of Neuroscience Methods, 195(1):92–106, 2011. ISSN 01650270. doi: 10.1016/j.jneumeth.2010.11. 020.

24. Thomas Kreuz, Daniel Chicharro, Conor Houghton, Ralph G Andrzejak, and Florian Mormann. Monitoring spike train synchrony. Journal of Neurophysiology, 109(5):1457–1472, mar 2013. ISSN 0022-3077. doi: 10.1152/jn.00873.2012.

25. Mario Mulansky and Thomas Kreuz. PySpike—A Python library for analyzing spike train synchrony. SoftwareX, 5:183–189, 2016. ISSN 23527110. doi: 10.1016/j.softx.2016.07.006.

26. Eric Jones, Travis Oliphant, Pearu Peterson, et al. SciPy: Open source scientific tools for Python, 2001–. [Online; accessed 27 August 2018].

27. Peter Dayan and L F Abbott. Theoretical Neuroscience: Computational and Mathematical Modeling of Neural Systems. Computational Neuroscience. MIT Press, Cambridge, Massachusetts, 2001. ISBN 0-262-04199-5.

28. Peter Sterling and Simon Laughlin. Principles of Neural Design. MIT Press, Cambridge, Massachusetts, 2015. ISBN 9780262028707.

29. Jens-Oliver Muthmann, Hayder Amin, Evelyne Sernagor, Alessandro Maccione, Dagmara Panas, Luca Berdondini, Upinder S Bhalla, and Matthias Helge Hennig. Spike Detection for Large Neural Populations Using High Density Multielectrode Arrays. Frontiers in Neuroinformatics, 9(December):1–21, dec 2015. ISSN 1662-5196. doi: 10.3389/fninf.2015.00028.

30. Stephen M Carcieri, Adam L Jacobs, and Sheila Nirenberg. Classification of retinal ganglion cells: a statistical approach. Journal of neurophysiology, 90:1704–1713, 2003. ISSN 0022-3077. doi: 10.1152/jn.00127.2003.

31. Jonathan D Victor. Spike train metrics. Current Opinion in Neurobiology, 15(5):585–592, oct 2005. ISSN 09594388. doi: 10.1016/j.conb.2005.08.002.

32. Kevin Patrick Murphy. Machine Learning: A Probabilistic Perspective. Adaptive Computation and Machine Learning. MIT Press, 2012. ISBN 9780262018029.

33. Nguyen Xuan Vinh, Julien Epps, and James Bailey. Information theoretic measures for clusterings comparison: Variants, properties, normalization and correction for chance. The Journal of Machine Learning Research, 11:2837–2854, 2010. ISSN 15280020. doi: 10.1182/blood-2008-03-145946.

34. Robert Tibshirani, Guenther Walther, and Trevor Hastie. Estimating the number of clusters in a data set via the gap statistic. Journal of the Royal Statistical Society: Series B (Statistical Methodology), 63(2):411–423, 2001.

35. Ronen Segev, Jason Puchalla, and Michael J Berry. Functional organization of ganglion cells in the salamander retina. Journal of Neurophysiology, 95(4):2277–2292, 2006.

36. Sneha Ravi, Daniel Ahn, Martin Greschner, EJ Chichilnisky, and Greg D Field. Pathway-specific asymmetries between on and off visual signals. Journal of Neuroscience, pages 2008–18, 2018.

37. Olivier Marre, Dario Amodei, Nikhil Deshmukh, Kolia Sadeghi, Frederick Soo, Timothy E Holy, and Michael J Berry. Mapping a complete neural population in the retina. Journal of Neuroscience, 32(43):14859–14873, 2012.

38. Adam Bleckert, Gregory W Schwartz, Maxwell H Turner, Fred Rieke, and Rachel OL Wong. Visual space is represented by nonmatching topographies of distinct mouse retinal ganglion cell types. Current Biology, 24(3):310–315, 2014.

